# Chasing the origin of SARS-CoV-2 in Canada’s COVID-19 cases: A genomics study

**DOI:** 10.1101/2020.06.25.171744

**Authors:** Calvin P Sjaarda, Nazneen Rustom, David Huang, Santiago Perez-Patrigeon, Melissa L Hudson, Henry Wong, T Hugh Guan, Muhammad Ayub, Claudio N Soares, Robert I Colautti, Gerald A Evans, Prameet M Sheth

**Affiliations:** Queen’s Genomics Lab at Ongwanada (QGLO), Ongwanada Resource Center, Kingston, Ontario, Canada, K7M8A6; Department of Psychiatry, Queen’s University, Kingston, Ontario, Canada, K7L3N6; Centre for Neuroscience Studies, Queen’s University, Kingston, Ontario Canada, K7L3N6; Biology Department, Queen’s University, Kingston, Ontario, Canada; Department of Medicine, Division of Infectious Diseases, Queen’s University, Kingston, Ontario; Division of Microbiology, Kingston Health Sciences Center, Kingston, Ontario, Canada; Department of Family Medicine, Queen’s University, Kingston, Ontario, Canada; Department of Pathology and Molecular Medicine, Queen’s University, Kingston, Ontario, Canada; Department of Biomedical & Molecular Sciences, Queen’s University, Kingston, Ontario; Gastrointestinal Disease Research Unit, Kingston Health Sciences Center, Kingston, Ontario

## Abstract

The emergence and global spread of SARS-CoV-2 has had profound social and economic consequences and has shed light on the importance of continued and additional investment in global health and infectious disease surveillance. Identifying changes in viral genomes provides key insights into viral diversity, how viruses spread within populations, and viral strategies for evasion of host immune systems. Here we report twenty-five SARS-CoV-2 genome sequences collected from some of the first COVID-19 cases in eastern Ontario, Canada (March 18-30, 2020). The reported genomes belong to the S-clade (*n*=2) and G-clade (*n*=23) of SARS-CoV-2 and contain 45 polymorphic sites including one shared missense and three unique synonymous variants in the gene encoding the spike protein. A phylogenetic analysis enabled the tracing of viral origin and potential transmission into and within Canada. There may be as many as sixteen unique infection events represented in these samples, including at least three that were likely introduced from Europe and seven from the USA. In addition, four separate genomes are each shared by multiple patients, suggesting a common origin or community spread even during this early stage of infection. These results demonstrate how molecular epidemiology and evolutionary phylogenetics can help local health units track origins and vectors of spread for emerging diseases like SARS-CoV-2. Earlier detection and screening in this way could improve the effectiveness of regional public health interventions to prevent future pandemics.

## Introduction

The past two decades have seen the emergence of three novel betacoronaviruses that have been associated with outbreaks in the human population including Severe Acute Respiratory Syndrome CoronaVirus (SARS-CoV) in 2002^1^, Middle East Respiratory Syndrome (MERS-CoV) in 2012^2^, and SARS-CoV-2 in 2019^3,4^. All three coronaviruses appear to have originated from bats and likely were transmitted to humans by zoonotic transmission possibly through an intermediate vertebrate vector^5,6^.

COrona VIrus Disease – 2019 (COVID-19) is the infectious disease caused by the SARS-CoV-2. The first confirmed case dates to December 8, 2019 in a patient from Wuhan City in the Hubei province of China^7^. The virus quickly spread through Wuhan and neighbouring parts of Hubei province despite rapid and aggressive public health interventions^8^. The following months led to global spread of the virus and was officially classified as a pandemic by the WHO on March 11, 2020. The spread of SARS-CoV-2 has had devastating consequence to human health with over 9.12 million documented cases and 473,797 deaths as of June 24, 2020 including 101,637 confirmed cases and 8436 deaths in Canada (WHO Situation Report Number 156).

SARS-CoV-2 is a spherical, enveloped particle, positive-sense, single stranded RNA genome that is 29.9 kb in length^3,9^. Genome organization of SARS-CoV-2 has the characteristic gene order 5’-replicase ORF1ab, spike (S), envelope (E), membrane (M), and nucleocapsid (N)-3’^6^. Coronavirus S proteins bind the host receptor enabling viral entry to the cell and have demonstrated the highest sequence variability in the viral genome^10^. The S protein in both SARS-CoV and SARS-CoV-2 interact with the host’s angiotensin-converting enzyme 2 (ACE2)^11^; however, the spike protein in SARS-CoV-2 has ∼10- to 20-fold higher binding affinity than SARS-CoV^12^.

The complete SARS-CoV-2 genome was published on Jan 12, 2020 from a patient in Wuhan, China^3^. The collaborative effort of many labs around the world have now published the genomes of many SARS-CoV-2 genomes including 53,992 in GISAID (www.gisaid.org), 24,438 in NCBI (www.ncbi.nlm.nih.gov), and 7,989 in ViPR (www.viprbrc.org) (as of June 24, 2020). Allelic differences with biased geographic distribution have been observed between three major SARS-CoV-2 clades, likely the consequence of founder effects^13^. Superclade I (a superset of V-clade defined in GISAID EpiCoV portal^14^), has limited variability with respect to the reference genome and has been described in the majority of isolates in Asia^13^. Superclade II (S-clade) has two characteristic variants at C3037T and T28144C^13^ and includes most viral isolates from the USA, especially the West Coast^13,15^. Superclade III, (G-clade) has four characteristic variants at C241T, C3037T, C14408T, and A23403G^13^, and includes most viral isolates from Europe and many on the East Coast of the USA^13,15^. The A23403G mutation results in a D614G substitution in the S protein which may result in increased fitness as this viral strain is rapidly and repeatedly replacing other forms of the virus around the globe^16^. This mutation is also embedded in an immunological epitope which elicited antibody production in patients during the 2003 SARS-CoV epidemic^17^, and may mediate antibody escape, making individuals susceptible to a second infection of COVID-19^16^.

Databases containing tens of thousands of SARS-CoV-2 genomes provide an unprecedented opportunity to reconstruct the establishment and spread of the virus in specific locales. We demonstrate the ability to trace the origin and community spread of SARS-CoV-2 using sequence data from twenty-five viral genomes isolated from some of first cases of COVID-19 in the eastern region of the province of Ontario, Canada. This knowledge may improve the effectiveness of public health interventions to prevent future pandemics

## Methods

### Sample collection

Nasopharyngeal (NP) swabs were collected in viral transport media from symptomatic patients being tested for SARS-CoV-2 at Kingston Health Sciences Center (KHSC) and the surrounding hospitals. Extraction of total RNA from viral transport media was performed using the Maxwell RSC Whole blood RNA/DNA kit (Promega Corporation, Madison, WI) on the Maxwell RSC 16 automated nucleic acid extractor. Each sample was tested for the presence of SARS-CoV-2 using a laboratory developed multiplex real-time PCR assay targeting the Envelope (E) and RNA dependent RNA polymerase (RdRp) genes^18^ on the ViiA7 Real-Time PCR System.

### SARS-CoV-2 genome sequencing

The extracted nucleic acids from COVID-19 positive cases were anonymized and shared with Queen’s Genomics Lab at Ongwanada (Q-GLO). RNA from each sample (5 μl) was reverse transcribed to complimentary DNA using the SuperScript VILO cDNA Synthesis Kit on a SimpliAmp Thermal Cycler. Libraries were constructed manually using the Ion AmpliSeq SARS-CoV-2 Research Panel, Ion Xpress Barcodes, and the Ion AmpliSeq Library Kit Plus following the manufacturer’s recommendations including amplification cycles based on viral load. Templating and chip loading were performed on the Ion Chef system using the Ion 510 & Ion 520 & Ion 530 Kit-Chef. Up to sixteen samples were multiplexed on an Ion 530 chip and sequenced using the Ion GeneStudio S5 Plus Semiconductor Sequencer.

### Chart review

Demographic data and travel history were extracted from a review of laboratory requisitions, hospital charts, and public health case investigation charts. Assessment of linked cases were based off public health case investigation charts.

### Data analysis

Preliminary analysis was performed on a Torrent Suite Server and using custom plug-ins created by ThermoFisher Scientific specifically for the Ion Ampliseq SARS CoV-2 panel including AssemblerTrinity for genome-guided assembly of the viral genome, IRMAreport to build a consensus sequence, and COVID19AnnotateSnpEff to annotate variants. VCF files generated by Torrent Suite Variant Caller were filtered to remove eleven variants with read depth less than 1000 and quality score less than 400. Filtered variants were used for sample clustering with the Maximum Likelihood Tree^19^ in Molecular Evolutionary Genetics Analysis (MEGA) software^20,21^.

### Phylogenetic analysis

A data analysis pipeline using molecular phylogenetics was developed to reconstruct infection origins and spread utilizing two main assumptions. First, identical sequences are identical by descent – that is, the same variants are unlikely to have evolved independently. Second, new mutations accrue at an average rate of 1 base pair (bp) every 7 to 21 days. This is based on an average of 24.225 bp substitutions per year in the Nextstrain analysis^22^ of the GISAID^14,23^ database. Briefly, the viral genome sequenced from each patient was used as the query sequence in a BLAST+^24^ search against a local reference containing 25,132 SARS Cov-2 genomes from the GISAID database (accessed May 15, 2020). Only highly similar sequences with no more than two mismatches were retained from the database reducing the number of reference genomes from 25,132 to 1,251. Many references were identical genomes isolated in separate individuals, which were collapsed to 72 unique reference sequences. A reference sequence from Wuhan (Wuhan-Hu1/2019) was added to root the phylogenetic tree. Fifty-nine of these 72 reference sequences shared the same common ancestor but differed by one or two substitutions, or otherwise formed derived groups that were not informative to reconstructing the ancestry of the sample genomes. These extraneous tips were removed, leaving only the patient-derived genomes and 13 unique and informative ancestral sequences. Phylogenies were built using the ape (v5.3) package in R, using the maximum likelihood approach implemented in the *optim.pml* function with 100 bootstrap iterations. The ggtree (v2.0.1) and ggplot2 (v3.2.1) packages in R were used to generate tree visualizations.

## Results

Sample demographics including age and biological sex are shown in Table 1. At time of analysis, thirty-two SARS-CoV-2 samples from local cases of COVID-19 were made available, however seven samples did not pass quality control (insufficient viral load or low sequencing coverage) and were excluded from further analysis. For the remaining twenty-five samples, an average of 1.3 million mapped reads per sample was generated, with an average of 8,175 mean read depth and 96.06% uniform coverage (Supplementary Table 1). Twenty-one samples have 100% coverage of the genome and four samples have greater than 99.4% coverage. Samples 19 and 21 had the lowest uniformity in coverage which resulted in several gaps in the consensus sequence, so these two sequences were removed from phylogenetic analysis. Most viral genomes contained between six to eight variants when compared with the NC_045512.2 reference genome with greater number of variants reported in Sample 41 (10 variants) and Sample 36 (12 variants). The most common nucleotide substitution is from cysteine to thymine (25/45 variants), followed by guanine to thymine (7/45 variants) and adenine to guanine (4/45). All the shared variants are homozygous variants except for heterozygous SNP in the orf1ab gene of Samples 11 and 30 (C9994A). We see an additional four heterozygous variants in three samples that are unique to that sample (Supplementary Table 2). Finally, we observe one shared missense variant (G25217T) and three unique synonymous variants (C24382T, T24982C, C25357T) in the gene encoding the S protein.

**Table 1.** Summary of patient demographic data including anonymous sample number, age, and biological sex of the participant (Male, Female), date of sample collection, and GISAID accession number for consensus sequence.

| Sample ID | Age | Sex | Sample collection date | Accession number in GISAID database |
| --- | --- | --- | --- | --- |
| 1 | 70s | F | March 18 | EPI_ISL_459866 |
| 2 | 40s | M | March 22 | EPI_ISL_459867 |
| 4 | 50s | F | March 24 | EPI_ISL_459868 |
| 10 | 20s | F | March 26 | EPI_ISL_459869 |
| 11 | 40s | F | March 26 | EPI_ISL_459871 |
| 12 | 70s | M | March 26 | EPI_ISL_459872 |
| 16 | 60s | M | March 26 | EPI_ISL_459873 |
| 17 | 50s | F | March 27 | EPI_ISL_459874 |
| 18 | 20s | M | March 27 | EPI_ISL_459875 |
| 19 | 70s | M | March 27 | EPI_ISL_459877 |
| 21 | 80s | M | March 27 | EPI_ISL_459878 |
| 23 | 20s | F | March 28 | EPI_ISL_459879 |
| 24 | 50s | M | March 28 | EPI_ISL_459880 |
| 25 | 50s | F | March 28 | EPI_ISL_459881 |
| 26 | 20s | M | March 28 | EPI_ISL_459882 |
| 29 | 60s | M | March 28 | EPI_ISL_459883 |
| 30 | 80s | F | March 28 | EPI_ISL_459884 |
| 34 | 80s | M | March 29 | EPI_ISL_459885 |
| 35 | 70s | M | March 30 | EPI_ISL_459886 |
| 36 | 40s | F | March 30 | EPI_ISL_459887 |
| 37 | 70s | M | March 30 | EPI_ISL_459888 |
| 38 | 20s | F | March 30 | EPI_ISL_459889 |
| 39 | 60s | M | March 29 | EPI_ISL_459890 |
| 41 | 70s | M | March 30 | EPI_ISL_459891 |
| 42 | 60s | F | March 30 | EPI_ISL_459892 |

**Table 2.** Summary of viral origin of first COVID-19 cases into the eastern region of the province of Ontario, Canada. Phylogenetic analysis determined viral origin based on ancestral sequences in the GISAID database and correlated with patient’s reported travel history. Samples with identical viral genome sequences are grouped into a single cluster and may represent shared origin or community transfer.

| Sample ID | Clade | Cluster | Origin based on phylogeny | Reported Travel History or contact with known case |
| --- | --- | --- | --- | --- |
| 23 | S-clade | 1 | North America – USA |  |
| 10 | G-clade 1 | 2 | Europe – UK/Spain | Travel: Europe – Ireland |
| 1 | G-clade 1 | 3 | Europe – UK/Portugal | Travel: Europe – Portugal |
| 12 | G-clade 1 | 4 | Europe – UK/Spain | Travel: Europe – Spain |
| 18 | G-clade 2 | 5 | North America – USA |  |
| 17 | G-clade 2 | 6 | North America – USA |  |
| 38 | G-clade 2 | 6 | North America – USA |  |
| 39 | G-clade 2 | 6 | North America – USA |  |
| 42 | G-clade 2 | 6 | North America – USA |  |
| 24 | G-clade 2 | 7 | North America – USA | No travel: contact Samples 25, 26 |
| 25 | G-clade 2 | 7 | North America – USA | No travel: contact Samples 24, 26 |
| 26 | G-clade 2 | 7 | North America – USA | No travel: contact Samples 24, 25 |
| 37 | G-clade 2 | 8 | North America – USA |  |
| 2 | G-clade 2 | 9 | North America – USA | Travel: North America – Barbados |
| 16 | G-clade 2 | 10 | North America – USA | No travel: contact Sample 4 |
| 4 | G-clade 2 | 11 | North America – USA | No travel: contact Sample 16 |
| 34 | G-clade 2 | 11 | North America – USA | Travel: USA – Florida |
| 35 | G-clade 2 | 12 | North America – USA |  |
| 36 | G-clade 2 | 13 | North America – USA |  |
| 29 | G-clade 2 | 14 | North America – USA |  |
| 11 | G-clade 2 | 15 | North America – USA | No travel: contact out of province |
| 30 | G-clade 2 | 15 | North America – USA |  |
| 41 | G-clade 2 | 16 | North America – USA | Travel: USA – Arizona |

Clustering of the 25 viral sequences within the maximum likelihood tree (Fig 1A) is driven by variants that are shared between samples (Fig 1B) and branch length is driven by unique variants carried by each sample (Supplementary Table 2). The samples formed two large clades including two samples that carry two mutations that correspond to the S-clade (C8782T and T28144C) and twenty-three samples that carry four mutations reported in the G-clade (C241T, C3037T, C14408T, and A23403G). Within the G-clade we see smaller clusters that correspond to cluster 2 (Samples 17, 18, 24, 25, 26, 37, 38, 39, and 42), cluster 5 (Samples 1, 10 and 12), and cluster 9 (Samples 2, 4, 11, 16, 29, 30, 34, 35, 36, and 41) reported by Korber et al.^16^.

**Figure 1.**
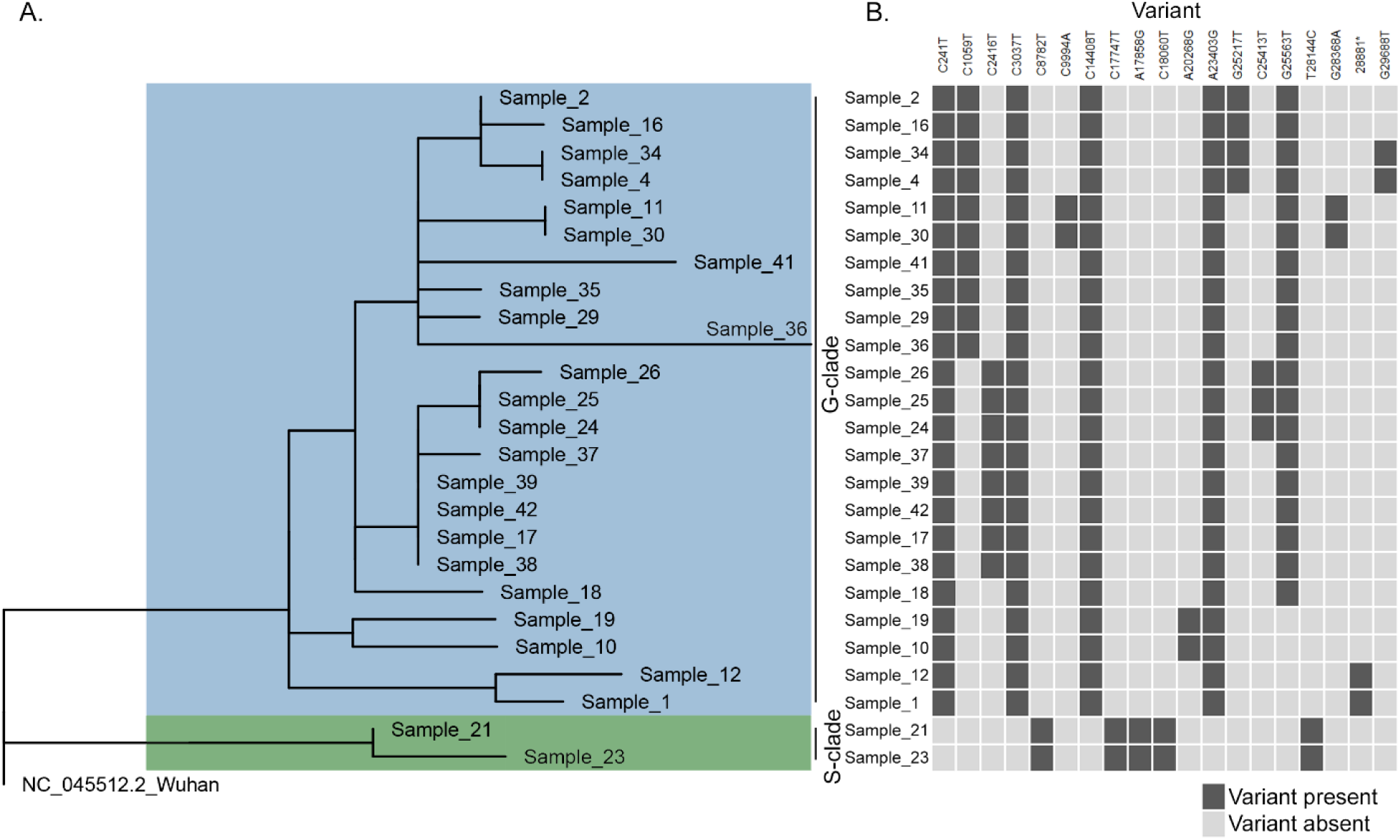
Clustering based on 25 SARS-CoV-2 genome sequences isolated from the early cases of COVID-19 in eastern Ontario. A. Maximum Likelihood Tree, rooted on the Wuhan reference strain, conducted by MEGA X using 45 variants discovered in 25 samples. B. Distribution of variants shared by two or more samples. Twenty-three viruses carry variants that characterize the G-clade of SARS-CoV-2 and two carry variants of the S-clade. *Variant at position 28881 is a GGG → AAC substitution.

Phylogenetic analysis using local patient SARS-CoV-2 genomes were matched with BLAST+ to reference sequences published in the GISAID database. This resulted in 13 ancestral genomes relevant to retracing the origins and spread of local COVID-19 cases (Fig 2). Sample 23 belongs to the S-clade of SARS-CoV-2 and is derived from the ancestral genome shared by reference 1. Samples sharing this genome are almost exclusively from North America and predominately from the USA (Supplementary Table 3). Samples 1, 10, and 12 belong to the G-clade cluster 1. This clade contains ancestral reference genomes (16, 39, 46, 68, 69, and 70) reported all over the world, but over-represented in Europe, especially the UK and Spain. Reported travel history supports European origin in these three samples (Table 2). The other nineteen samples in the G-clade cluster 2 are derived from several ancestral sequences (3, 14, 53, 57, 67, 72) which are predominantly from North American (USA) patients. Reported travel history supports an American origin of infection for patients 34 and 41, who reported visiting the USA before being diagnosed with the virus (Table 2). Within the G-clade 2 samples, we observed four clusters of two or more samples with the same viral genome indicating a common source of infection (Fig 2; Table 2).

**Figure 2:**
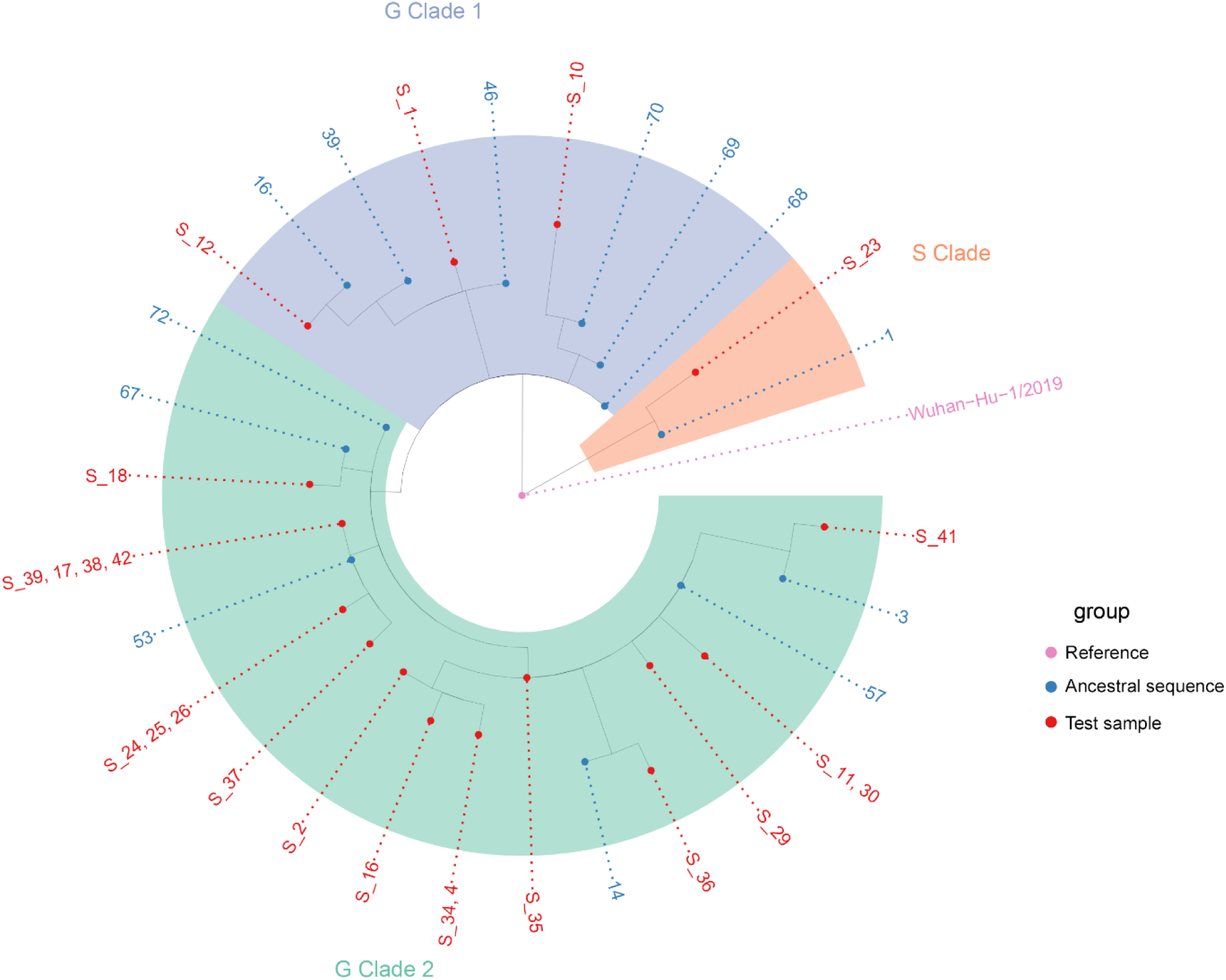
Nextstrain phylogenetic tree of local cases of SARS-CoV-2 and the most similar ancestral sequences in the GISAID database. Sample 23 in the S-clade appears derived from ancestral sequences 1 which predominately includes sequences from North America (USA). Samples 1,10, and 12 in the G-clade 1 is derived from ancestral sequence 68 and others which predominately appear European (UK, Spain, Portugal) in origin. The other 19 samples in the G-clade 2 is derived from ancestral sequence 72 and others which appear predominantly North American (USA) in origin.

## Discussion

We analyzed twenty-five SARS-CoV-2 genomes from cases of COVID-19 within the eastern region of the province of Ontario, Canada. These samples included most of the first cases in the region and therefore infection was assumed to have occurred during international travel or while in proximity to an individual who had recently travelled. Genome variation among these samples has important implications for the pathology and epidemiology of this disease, as discussed below.

A major concern with novel viruses is mutation rates and how novel mutations will affect virulence, vaccine development, and reinfection^15-17,25,26^. Of primary significance are mutations in the S protein because the spike protein defines viral host range and is often the target of neutralizing antibodies^27,28^. This project identified three novel unique mutations in the coding region of the S protein, however all three are predicted to be synonymous and likely will not affect viral virulence or epitopes. A fourth polymorphic site at C25217T occurs in four samples (2, 4, 34, and 16) is a missense variant resulting in a glycine to cysteine substitution at the 1219^th^ residue of the S protein. In silico modeling or functional validation studies may describe the impact of this mutation on the function of the transmembrane domain. This project identified five heterozygous variants C360T, C9729A, G10818T, G17325T, C18136T. Four of these heterozygous variants were unique to one sample and had alternate allele frequencies ranging from 25-39% suggesting an emerging viral mutation that occurred since infection of the host. However, one heterozygous variant, C9729A, was shared between two samples (Samples 11 and 30) and had an alternate allele frequency of 0.58 and 0.59 indicating that these two individuals likely have superinfection of two SARS-CoV-2 strains^29^.

Clustering of shared mutations identified two samples (21 and 23) that belong to the S-clade of virus and characterized by two polymorphic sites at C8782T and C28144T^13^. The presence of three additional polymorphic sites at C17747T, A17858G, and C18060T exclusively present in North America^26^ support a North American origin for these two samples. Furthermore, phylogenetic analysis identified potential ancestral samples for Sample 23 that were predominately isolated in the USA. The other twenty-three samples share four shared polymorphic sites at C3037T, C14408T, A23404G, and A23403G and belong to the G-clade of viruses^13^. Within the G-clade we observe two distinct clusters, G-clade 1 which appears European in origin and G-clade 2 which appears to originate in the USA. Reported travel history supports our phylogenetics analysis for European origin for the G-clade 1 as Sample 1 reported recent travel to Portugal, Sample 10 travelled to Ireland, and Sample 12 to Spain. Similarly, reported travel history supports North American origin for the G-clade 2 as Sample 34 reported recent travel to Florida, USA and Sample 41 reported recent travel to Arizona, USA. These correlations between phylogenetic origin and reported travel history indicates how viral genome sequencing can successfully trace the origin of SARS-CoV-2 infections into Canada. Interestingly, we observed several clusters of samples that shared the same viral sequence indicating that these samples either had a shared source of infection or a result of community transfer. Chart review demonstrated that Samples 24, 25, and 26 did not travel outside of Canada, they are contacts of each other, and share the same SARS-CoV-2 strain, demonstrating community transfer in eastern Ontario was likely even during the early stages of infection. Similarly, samples 17, 38, 39 and 42 all share the same viral strain but the connection between these individuals is unknown. Identification of the common thread among these patients (or other clusters of cases) could help to identify major sources/pathways of infection. Infection in Sample 34 is likely attributed to their travel to the USA (Table 2); however, Samples 4 and 34 share the same viral strain and Sample 4 tested positive for COVID-19 five days earlier than Sample 34. A connection between Sample 4 and 34 is unknown, as are the travel details of Sample 34, but these observations raise the possibility that infection in Sample 34 may not be related to their travel to the USA. These results demonstrate how sample clustering and phylogenetic analysis based on viral genome sequencing may assist and prioritize case and contact tracing to further monitor and contain the spread of the virus.

There were several limitations in using genomics to trace viral origin and path of transmission of SARS-CoV-2. First, in the early stages of the pandemic, most countries were only screening and testing international travelers who displayed symptoms. This allowed asymptomatic carriers to go undetected masking potential sources of infection. As a result, gaps exist in the databases reporting SARS-CoV-2 genomes masking the true sources of infection. However, the rapid publication of genome sequences from around the world can help to offset this limitation, by identifying geographical clusters and specific genomic variants that are shared across regions. The relatively high mutation rate helps to distinguish primary and secondary infections in the span of one to two weeks. A second limitation is a lack of detailed travel and interaction histories for patients due to differences in reporting and data collection among collection sites and agencies. Rigorous adherence to standardized data collection protocols, like WHO’s guidance for contact tracing in the context of COVID-19 coupled with genomics data as described here, may facilitate effective contact tracing that is required to break the chains of viral transmission. A final limitation is that sequencing SARS-CoV-2 in COVID-19 cases with low viral load was problematic due to lack of RNA input for library construction as seen by the number of excluded samples and those with low coverage uniformity (Samples 19 and 21). However, this will become less limiting over time as high-throughput sequencing devices continue to improve in sensitivity and throughput.

In summary, this is the first description of SARS-CoV-2 genomes in COVID-19 positive cases in eastern Ontario. These are many of the first detected cases in eastern Ontario and infection was believed to be foreign in origin and the result of international travel or proximity to an individual who had travelled. This analysis illustrates that many of the infections originated from our geographical neighbour, the USA, but other sources of infection include several countries in Europe. We also observed community transfer in cases that did not travel out of the country. These results demonstrate how molecular epidemiology and evolutionary phylogenetics can help local health units to track origins and vectors of spread for emerging diseases like SARS-CoV-2. Earlier detection and screening and alternative modes for contact tracing may improve the effectiveness of regional public health interventions to prevent future pandemics.

## Acknowledgements

We acknowledge the authors from the originating laboratories responsible for obtaining the specimens and the submitting laboratories where genetic sequence data were generated and shared via the GISAID Initiative. We gratefully acknowledge Ongwanada Resource Center and its board of directors for their support of Q-GLO and this project.

## Author Contributions

CPS, NR, and PMS conceived and designed the study. GAE, SPP, and THG were involved in patient care and chart review, HW and PMS performed collection and testing of biological materials. CPS performed the sequencing experiments and preliminary data analysis. DH and RIC performed the phylogenetic analyses. CPS drafted the manuscript. All authors contributed to manuscript revisions and approved the submitted version.

## Ethics Statement

Biological samples and demographic data were collected from patients within the circle of care resulting from clinical testing for SARS-CoV-2. Samples were anonymized and de-identified so researchers were blind to the identity of the patients. Only secondary non-identifying data including age, biological sex, and travel history were provided. All experimental protocols were approved by and conducted in accordance with the Queen’s University Health Sciences and Affiliated Teaching Hospitals Research Ethics Board (PSIY-676-20).

## Data Availability Statement

The consensus sequence for each sample was submitted to GISAID (https://www.gisaid.org/) under the accession IDs provided in Table 1.

## References

1 Lee, N. et al. A Major Outbreak of Severe Acute Respiratory Syndrome in Hong Kong. New England Journal of Medicine 348, 1986–1994, doi:10.1056/NEJMoa030685 (2003).

2 Zaki, A. M., van Boheemen, S., Bestebroer, T. M., Osterhaus, A. D. M. E. & Fouchier, R. A. M. Isolation of a Novel Coronavirus from a Man with Pneumonia in Saudi Arabia. New England Journal of Medicine 367, 1814–1820, doi:10.1056/NEJMoa1211721 (2012).

3 Wu, F. et al. A new coronavirus associated with human respiratory disease in China. Nature 579, 265–269, doi:10.1038/s41586-020-2008-3 (2020).

4 Gorbalenya, A. E. et al. The species Severe acute respiratory syndrome-related coronavirus: classifying 2019-nCoV and naming it SARS-CoV-2. Nature Microbiology 5, 536–544, doi:10.1038/s41564-020-0695-z (2020).

5 Cui, J., Li, F. & Shi, Z.-L. Origin and evolution of pathogenic coronaviruses. Nature Reviews Microbiology 17, 181–192, doi:10.1038/s41579-018-0118-9 (2019).

6 Lu, R. et al. Genomic characterisation and epidemiology of 2019 novel coronavirus: implications for virus origins and receptor binding. The Lancet 395, 565–574, doi: https://doi.org/10.1016/S0140-6736(20)30251-8 (2020).

7 Wu, Z. & McGoogan, J. M. Characteristics of and Important Lessons From the Coronavirus Disease 2019 (COVID-19) Outbreak in China: Summary of a Report of 72314 Cases From the Chinese Center for Disease Control and Prevention. JAMA, doi:10.1001/jama.2020.2648 (2020).

8 Tang, X. et al. On the origin and continuing evolution of SARS-CoV-2. National Science Review, doi:10.1093/nsr/nwaa036 (2020).

9 Mousavizadeh, L. & Ghasemi, S. Genotype and phenotype of COVID-19: Their roles in pathogenesis. Journal of Microbiology, Immunology and Infection, doi: https://doi.org/10.1016/j.jmii.2020.03.022 (2020).

10 Woo, P. C. Y., Huang, Y., Lau, S. K. P. & Yuen, K.-Y. Coronavirus genomics and bioinformatics analysis. Viruses 2, 1804–1820, doi:10.3390/v2081803 (2010).

11 Walls, A. C. et al. Structure, Function, and Antigenicity of the SARS-CoV-2 Spike Glycoprotein. Cell 181, 281-292.e286, doi: https://doi.org/10.1016/j.cell.2020.02.058 (2020).

12 Wrapp, D. et al. Cryo-EM structure of the 2019-nCoV spike in the prefusion conformation. Science 367, 1260–1263, doi:10.1126/science.abb2507 (2020).

13 Chiara, M., Horner, D. S., Gissi, C. & Pesole, G. Comparative genomics suggests limited variability and similar evolutionary patterns between major clades of SARS-CoV-2. bioRxiv, 2020.2003.2030.016790, doi:10.1101/2020.03.30.016790 (2020).

14 Shu, Y. & McCauley, J. GISAID: Global initiative on sharing all influenza data - from vision to reality. Euro Surveill 22, 30494, doi:10.2807/1560-7917.ES.2017.22.13.30494 (2017).

15 Brufsky, A. Distinct Viral Clades of SARS-CoV-2: Implications for Modeling of Viral Spread. Journal of Medical Virology n/a, doi:10.1002/jmv.25902.

16 Korber, B. et al. Spike mutation pipeline reveals the emergence of a more transmissible form of SARS-CoV-2. bioRxiv, 2020.2004.2029.069054, doi:10.1101/2020.04.29.069054 (2020).

17 Wang, Q. et al. Immunodominant SARS Coronavirus Epitopes in Humans Elicited both Enhancing and Neutralizing Effects on Infection in Non-human Primates. ACS Infect Dis 2, 361–376, doi:10.1021/acsinfecdis.6b00006 (2016).

18 Corman, V. M. et al. Detection of 2019 novel coronavirus (2019-nCoV) by real-time RT-PCR. Euro Surveill 25, doi:10.2807/1560-7917.Es.2020.25.3.2000045 (2020).

19 Tamura, K. & Nei, M. Estimation of the number of nucleotide substitutions in the control region of mitochondrial DNA in humans and chimpanzees. Molecular Biology and Evolution 10, 512–526, doi:10.1093/oxfordjournals.molbev.a040023 (1993).

20 Stecher, G., Tamura, K. & Kumar, S. Molecular Evolutionary Genetics Analysis (MEGA) for macOS. Molecular Biology and Evolution 37, 1237–1239, doi:10.1093/molbev/msz312 (2020).

21 Kumar, S., Stecher, G., Li, M., Knyaz, C. & Tamura, K. MEGA X: Molecular Evolutionary Genetics Analysis across Computing Platforms. Molecular biology and evolution 35, 1547–1549, doi:10.1093/molbev/msy096 (2018).

22 Hadfield, J. et al. Nextstrain: real-time tracking of pathogen evolution. Bioinformatics 34, 4121–4123, doi:10.1093/bioinformatics/bty407 (2018).

23 Elbe, S. & Buckland-Merrett, G. Data, disease and diplomacy: GISAID’s innovative contribution to global health. Glob Chall 1, 33–46, doi:10.1002/gch2.1018 (2017).

24 Camacho, C. et al. BLAST+: architecture and applications. BMC Bioinformatics 10, 421, doi:10.1186/1471-2105-10-421 (2009).

25 Sanjuán, R., Nebot, M. R., Chirico, N., Mansky, L. M. & Belshaw, R. Viral Mutation Rates. Journal of Virology 84, 9733–9748, doi:10.1128/jvi.00694-10 (2010).

26 Pachetti, M. et al. Emerging SARS-CoV-2 mutation hot spots include a novel RNA-dependent-RNA polymerase variant. Journal of Translational Medicine 18, 179, doi:10.1186/s12967-020-02344-6 (2020).

27 Chen, W.-H., Hotez, P. J. & Bottazzi, M. E. Potential for developing a SARS-CoV receptor-binding domain (RBD) recombinant protein as a heterologous human vaccine against coronavirus infectious disease (COVID)-19. Human Vaccines & Immunotherapeutics, 1–4, doi:10.1080/21645515.2020.1740560 (2020).

28 Yuan, M. et al. A highly conserved cryptic epitope in the receptor binding domains of SARS-CoV-2 and SARS-CoV. Science 368, 630–633, doi:10.1126/science.abb7269 (2020).

29 Wen, S. & Zhang, X. A High-Coverage SARS-CoV-2 Genome Sequence Acquired by Target Capture Sequencing. medRxiv, 2020.2004.2011.20061507, doi:10.1101/2020.04.11.20061507 (2020).

