## Supplemental Data for "Chasing the origin of SARS-CoV-2 in Canada’s COVID-19 cases: A genomics study"

**Affiliations:**

*Observations on ancestral sequences (Fig 2; Supplementary Table 3)*
S-clade: Ancestral sequence (A) 1 is the ancestor of the Sample (S) 23 in the S-Clade. S23 has 2bp changes from A1, suggesting that there is a secondary source of infection from A1 to S23 that may not be represented in the database. Based on the reported mutation rate, S23 was infected ~3-5 weeks after the group in A1.

G-clade 1: A68 is ancestral to all infections in the G-clade 1. Individuals in A69 were infected ~1-3 weeks after A68, followed by A70 after another ~1-3 weeks. S10 is indirectly a result of the A70 infection but the S10 genome contains 3 unique bp mutations so there is a ~5-7 week window of potential transmission between A70 and S10.
A46 infections occurred about 5-7 weeks from an ancestral genome in group A68. S1 is derived from A46 but shows a 1bp mutation suggesting a ~1-3 week window of transmission. S12 is also derived from A46 but there is an identical sequence in Nextstrain represented by A16 reported in Canada. Since S12 reported travel to Spain, the A16 sequence reported in Canada may be a parallel infection from a similar ancestral host.

G-clade 2: A72 is the basal infection of G-clade 2, leading to ancestral strains A53, A67 and A57 within ~1-3 weeks. A3 and A14 are also derived from A72 but with a delay of about 5-7 weeks. S18 differs from A67 from a single bp suggesting infection ~1-3 weeks after A67. S17, S38, S39, S42 all have the same genotype as A53, suggesting a common origin for each infection. S24, S25, S26 have the same genotype but differ from A53 by about a week while S37 has a different mutation from A53 with the same divergence time. Thus S24, S25 and S26 appear to share the same source whereas S37 has a different source. However, these two sources were both derived from A53 by ~1-3 weeks. This suggests a potentially interesting chain of infections: A72 → A53/S17/S38/S39 → S24/S25/S26 as a group, and separately S37.
S35 matches A57 suggesting they share the same origin. S2, S4, S11, S16, S29, S30, and S34 are all derived from A57/S35 with a lag of ~1-3 weeks (S2 and S29) or 3-5 weeks. S11 and S30 are identical and share a common origin. S4 and S34 are also identical to each other and share an origin. S4, S34 and S16 each have 1bp mutation difference from S2 suggesting that S2 is closer to the origin of infection. These observations suggest a potentially interesting chain of infections: A57/S35 → S29, S11/S30, and S2 as 3 separate pathways, then S2 → S16 and S4/S34 as 2 separate pathways. S41 and S36 are both derived from the same source (A57/S35) but likely via separate pathways over a ~5-7 week time frame. The closest source to S41 is group A3 and the closest to S36 is group A14.

Table 1. Summary of viral genome sequencing for each sample.

| Sample ID | Number of mapped reads | Mean read depth | Coverage uniformity | Number of variants |
|---|---|---|---|---|
| 1 | 1,582,943 | 9,215 | 97.80% | 6 |
| 2 | 1,429,307 | 9,478 | 96.47% | 7 |
| 4 | 1,730,539 | 11,809 | 97.89% | 8 |
| 10 | 1,429,167 | 8,851 | 99.69% | 7 |
| 11 | 639,006 | 4,305 | 97.58% | 8 |
| 12 | 1,148,216 | 7,560 | 98.21% | 7 |
| 16 | 1,368,607 | 9,294 | 98.00% | 8 |
| 17 | 1,573,637 | 10,745 | 98.13% | 6 |
| 18 | 1,316,806 | 6,920 | 99.17% | 7 |
| 19 | 1,044,713 | 3,170 | 71.19% | 7 |
| 21 | 1,082,078 | 2,054 | 71.65% | 5 |
| 23 | 1,252,788 | 8,510 | 98.19% | 7 |
| 24 | 1,790,583 | 12,325 | 98.20% | 7 |
| 25 | 1,731,460 | 11,783 | 98.21% | 7 |
| 26 | 1,512,017 | 10,267 | 98.18% | 8 |
| 29 | 1,429,785 | 9,670 | 98.55% | 7 |
| 30 | 847,705 | 5,697 | 97.51% | 8 |
| 34 | 876,823 | 5,750 | 98.74% | 8 |
| 35 | 1,277,367 | 8,578 | 98.09% | 7 |
| 36 | 1,541,104 | 10,140 | 99.14% | 12 |
| 37 | 632,145 | 994 | 98.28% | 7 |
| 38 | 1,470,901 | 9,982 | 98.55% | 6 |
| 39 | 1,105,265 | 7,350 | 97.22% | 6 |
| 41 | 1,356,131 | 9,297 | 98.71% | 10 |
| 42 | 1,562,381 | 10,640 | 98.20% | 6 |

Table 2: Twenty-six unique variants were observed in thirteen SARS-CoV-2 viral genome sequences.

| Sample ID | Position | REF allele | ALT allele | Gene | HGVS_protein | Variant effect | Alternate allele frequency | Quality score | Read depth |
|---|---|---|---|---|---|---|---|---|---|
| 12 | 3373 | C | A | orf1ab | p.Asp1036Glu | missense | 1.000 | 2,977.9 | 6,129 |
| 23 | 4543 | C | T | orf1ab | p.Thr1426Thr | synonymous | 1.000 | 2,979.6 | 13,227 |
| 10 | 5230 | G | T | orf1ab | p.Lys1655Asn | missense | 1.000 | 2,981.8 | 5,175 |
| 16 | 6846 | T | A | orf1ab | p.Met2194Lys | missense | 1.000 | 2,982.2 | 5,740 |
| 12 | 9733 | C | T | orf1ab | p.Phe3156Phe | synonymous | 0.997 | 2,345.9 | 6,013 |
| 41 | 10188 | C | T | orf1ab | p.Thr3308Ile | missense | 0.998 | 2,946.2 | 5,429 |
| 37 | 10369 | C | T | orf1ab | p.Arg3368Arg | synonymous | 0.995 | 2,919.6 | 839 |
| 18 | 10507 | C | T | orf1ab | p.Asn3414Asn | synonymous | 1.000 | 2,976.7 | 6,788 |
| 35 | 11083 | G | T | orf1ab | p.Leu3606Phe | missense | 0.391 | 199.8 | 6,415 |
| 36 | 11916 | C | T | orf1ab | p.Ser3884Leu | missense | 0.995 | 2,912.3 | 8,359 |
| 36 | 12103 | A | G | orf1ab | p.Ser3946Ser | synonymous | 0.998 | 2,948.9 | 9,592 |
| 10 | 14481 | C | T | orf1ab | p.Thr4739Ile | missense | 0.993 | 1,026.0 | 141 |
| 19 | 17126 | T | C | orf1ab | p.Ser5621Pro | missense | 1.000 | 2,978.4 | 837 |
| 26 | 17590 | G | T | orf1ab | p.Leu5775Leu | synonymous | 0.298 | 83.5 | 6,791 |
| 36 | 18401 | C | T | orf1ab | p.Leu6046Leu | synonymous | 0.253 | 38.8 | 5,426 |
| 18 | 18877 | C | T | orf1ab | p.Val6204Val | synonymous | 1.000 | 2,964.7 | 7,981 |
| 36 | 18998 | C | T | orf1ab | p.His6245Tyr | missense | 0.998 | 2,948.9 | 14,265 |
| 1 | 19677 | G | T | orf1ab | p.Arg6471Met | missense | 1.000 | 2,974.8 | 2,630 |
| 41 | 24382 | C | T | S | p.Ser940Ser | synonymous | 0.998 | 2,948.9 | 4,962 |
| 41 | 24982 | T | C | S | p.Pro1140Pro | synonymous | 0.993 | 2,870.1 | 6,050 |
| 23 | 25357 | C | T | S | p.Leu1265Leu | synonymous | 0.998 | 2,952.1 | 2,181 |
| 29 | 27686 | C | T | ORF7a | p.Ser98Phe | missense | 1.000 | 2,972.1 | 12,615 |
| 19 | 27925 | C | T | ORF8 | p.Thr11Ile | missense | 0.973 | 2,699.8 | 5,540 |
| 41 | 27964 | C | T | ORF8 | p.Ser24Leu | missense | 1.000 | 2,964.9 | 29,390 |
| 36 | 28253 | C | T | ORF8 | p.Phe120Phe | synonymous | 0.290 | 74.9 | 15,904 |
| 36 | 29540 | G | A | ORF10 | . | upstream_gene | 0.995 | 2,919.7 | 4,006 |

Table 3: Location of published SARS-CoV-2 sequences predicted to be ancestral to viral genomes isolated for COVID-19 cases in the eastern region of the province of Ontario, Canada.

| Ancestral sequence ID | Region | Country | N |
|---|---|---|---|
| A_1 | North America | Canada | 2 |
| A_1 | North America | USA | 247 |
| A_1 | Oceania | Australia | 1 |
| A_3 | North America | USA | 6 |
| A_14 | Oceania | Australia | 1 |
| A_14 | North America | USA | 34 |
| A_14 | South America | Argentina | 1 |
| A_16 | North America | Canada | 2 |
| A_39 | Europe | Portugal | 4 |
| A_39 | Europe | Russia | 1 |
| A_46 | South America | Argentina | 1 |
| A_46 | Oceania | Australia | 15 |
| A_46 | Europe | Belgium | 17 |
| A_46 | North America | Canada | 4 |
| A_46 | Europe | Czech Republic | 1 |
| A_46 | Europe | Denmark | 8 |
| A_46 | Europe | United Kingdom | 101 |
| A_46 | Europe | France | 3 |
| A_46 | Asia | Georgia | 1 |
| A_46 | Europe | Germany | 6 |
| A_46 | Europe | Greece | 12 |
| A_46 | Europe | Hungary | 4 |
| A_46 | Europe | Iceland | 3 |
| A_46 | Asia | India | 1 |
| A_46 | Europe | Ireland | 1 |
| A_46 | Europe | Italy | 4 |
| A_46 | Asia | Japan | 5 |
| A_46 | Asia | Kuwait | 1 |
| A_46 | Europe | Latvia | 1 |
| A_46 | North America | Mexico | 1 |
| A_46 | Europe | Netherlands | 2 |
| A_46 | Oceania | New Zealand | 1 |
| A_46 | Europe | Portugal | 7 |
| A_46 | Europe | Russia | 18 |
| A_46 | Asia | Saudi Arabia | 2 |
| A_46 | Asia | Singapore | 1 |
| A_46 | Europe | Spain | 3 |
| A_46 | Asia | Sri Lanka | 1 |
| A_46 | Europe | Sweden | 4 |
| A_46 | Europe | Switzerland | 2 |
| A_46 | Asia | Taiwan | 2 |
| A_46 | Europe | Turkey | 1 |
| A_46 | Asia | United Arab Emirates | 1 |
| A_46 | North America | USA | 12 |
| A_46 | Asia | Vietnam | 7 |
| A_53 | Europe | Belgium | 1 |
| A_53 | North America | Canada | 1 |
| A_53 | Europe | United Kingdom | 1 |
| A_53 | Europe | France | 6 |
| A_53 | Europe | Russia | 1 |
| A_53 | Europe | Sweden | 2 |

| | | | |
|---|---|---|---|
| A_53 | North America | USA | 19 |
| A_57 | Oceania | Australia | 13 |
| A_57 | Europe | Austria | 9 |
| A_57 | South America | Brazil | 1 |
| A_57 | North America | Canada | 5 |
| A_57 | Europe | Denmark | 62 |
| A_57 | Europe | United Kingdom | 8 |
| A_57 | Europe | Finland | 2 |
| A_57 | Europe | France | 9 |
| A_57 | Europe | Germany | 15 |
| A_57 | Europe | Greece | 1 |
| A_57 | Asia | Israel | 2 |
| A_57 | Europe | Iceland | 28 |
| A_57 | Europe | Luxembourg | 8 |
| A_57 | Europe | Netherlands | 6 |
| A_57 | Europe | Norway | 3 |
| A_57 | Europe | Russia | 3 |
| A_57 | Asia | Singapore | 2 |
| A_57 | Europe | Spain | 1 |
| A_57 | Europe | Sweden | 1 |
| A_57 | Asia | Taiwan | 3 |
| A_57 | Asia | Thailand | 1 |
| A_57 | North America | USA | 278 |
| A_57 | Asia | United Arab Emirates | 3 |
| A_67 | Oceania | Australia | 2 |
| A_67 | North America | Canada | 6 |
| A_67 | Europe | Greece | 1 |
| A_67 | Asia | Japan | 3 |
| A_67 | Europe | Portugal | 1 |
| A_67 | Asia | Saudi Arabia | 1 |
| A_67 | Asia | Taiwan | 3 |
| A_67 | North America | USA | 33 |
| A_68 | Europe | Russia | 4 |
| A_68 | Asia | Saudi Arabia | 1 |
| A_68 | Europe | United Kingdom | 18 |
| A_68 | Asia | Singapore | 1 |
| A_68 | Europe | Sweden | 3 |
| A_68 | North America | USA | 21 |
| A_69 | South America | Argentina | 4 |
| A_69 | Oceania | Australia | 2 |
| A_69 | Europe | Denmark | 1 |
| A_69 | Europe | United Kingdom | 10 |
| A_69 | Asia | Georgia | 1 |
| A_69 | Europe | Greece | 1 |
| A_69 | Europe | Hungary | 4 |
| A_69 | Europe | Italy | 1 |
| A_69 | Europe | Luxembourg | 1 |
| A_69 | Europe | Netherlands | 3 |
| A_69 | Europe | Portugal | 1 |
| A_69 | Europe | Russia | 9 |
| A_69 | Europe | Spain | 18 |
| A_69 | Europe | Switzerland | 2 |
| A_69 | North America | USA | 4 |
| A_70 | Europe | United Kingdom | 1 |
| A_70 | Europe | Spain | 2 |
| A_72 | Europe | France | 2 |

| | | | |
|---|---|---|---|
| A_72 | Asia | Saudi Arabia | 1 |
| A_72 | North America | USA | 5 |
